# Latent Representations in Hippocampal Network Model Co-Evolve with Behavioral Exploration of Task Structure

**DOI:** 10.1101/2023.04.24.538070

**Authors:** Ian Cone, Claudia Clopath

## Abstract

Real-life behavioral tasks are often complex and depend on abstract combinations of sensory stimuli and internal logic. To successfully learn these tasks, animals must pair actions or decisions to the task’s complex structure. The hippocampus has been shown to contain fields which represent complex environmental and task variables, including place, lap, evidence accumulation, etc. Altogether, these representations have been hypothesized to form a “cognitive map” which encodes the complex real-world structure underlying behavior. However, it is still unclear how biophysical plasticity mechanisms at the single cell level can lead to the population-wide evolution of task-relevant maps. In this work we present a biophysically plausible model comprised of a recurrent hippocampal network and an action network, in which the latent representational structure co-evolves with behavior in a task-dependent manner. We demonstrate that the network develops latent structures that are needed for solving the task and does not integrate latent structures which do not support task learning. We show that, in agreement with experimental data, cue-dependent “splitters” can only be induced at the single cell level if the task requires a split representation to solve. Finally, our model makes specific predictions on how biases in behavior result from experimentally testable biases in the underlying latent representation.

## Introduction

Reinforcement learning algorithms, both artificial and biologically inspired, depend critically on the process described being Markovian – that is, actions and values can be assigned to given states (e.g. a place in the environment), irrespective of history^1–3^. Usually, the algorithm is concerned with learning a good policy (i.e. the strategy of the animal or agent) given an appropriate state space. Typically, in a simple 2D physical environment, an “appropriate state space” consists simply of locations within the environment. However, if an animal or an agent is learning a task which depends on previous history or abstract context not described by the state space (i.e., non-Markovian), simple tabular TD-learning (state value) or Q-learning (state-action value) will fail to find the solution. Therefore, one might also consider the problem of reinforcement learning in the inverse: what are the “appropriate” state representations upon which the policy can be described as Markovian, and how can we learn these representations^4,5^?

As an example of a non-Markovian task, imagine an agent starts at the top left of a 2D grid and traverses the space until it reaches a “reward port” at the bottom right of the grid. Upon reaching the port, the agent is only rewarded if it has previously traversed a “cue” location and is punished otherwise (**Figure 1a**). The optimal policy cannot be accurately described by single scalar values assigned to states (or state transitions) if they are defined as locations in this 2D space. This can be seen by examining the state-action values for a state immediately before the reward port (state 8 in the example of **Figure 1a**) – the transition from state 8 to the terminal state, state 9, is +1 if the agent has visited the cue state, but -1 if the agent has not. So, in Q-learning, Q(8,9) will not converge to the optimal policy. The most basic solution to this problem is to create two copies of that state (8’ and 8’’, in this example) before learning, using the external knowledge that the task depends on two cases; one where the agent has passed through the cue location, and one where the agent has not. However, this assumption breaks causality from the agent’s perspective. The only way to “know” if two copies of that state are required is to learn the task, but the only way to learn the task is to have two copies of that state. We are left with quite a conundrum.

**Figure 1.**
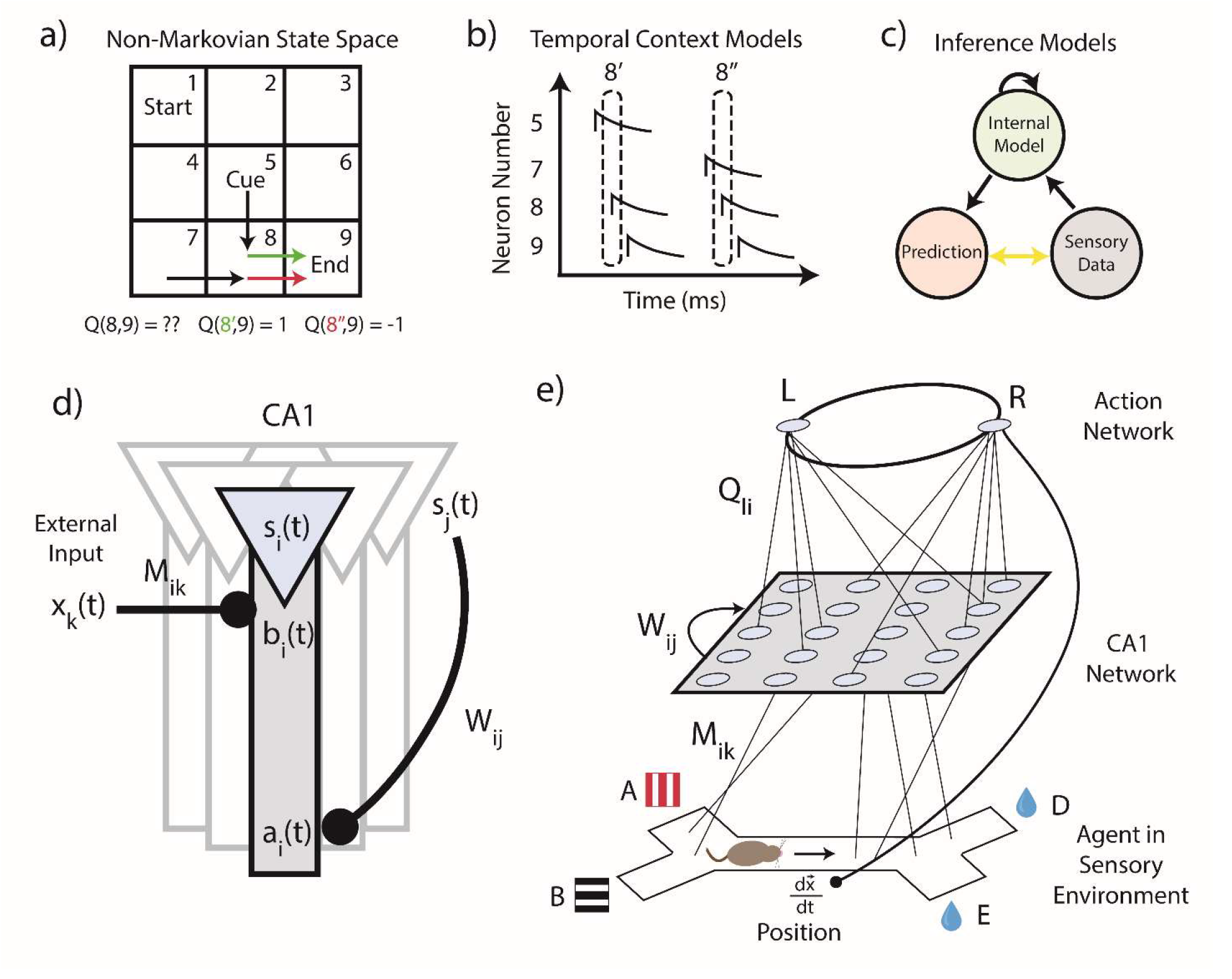
Feedback in three-compartment CA1 neurons Enables Context-Sensitive Representations. **a)** A 2D grid environment, in which the agent must visit a cue state (state 5) in order to receive a reward when it reaches the end state (state 9). Bottom, state transition 8-9 can be either rewarding (if preceded by 5) or punishing (if preceded by 7), leading to ambiguity in the value of the 8-9 transition. **b)** Some models include a history-dependence a priori as part of their state representation. Here, the population activity vector is equivalent to the state vector at that time, so the two potential “8” states (shown by the dotted ovals), are disambiguated. **c)** Alternatively, inference models compare internal predictions to external sensory observations, and update their internal models based on errors between their predictions and observations. Successfully trained inference models learn the latent structure of the task as part of their internal model. **d)** Schematic of the hippocampal network model we propose. The network receives external inputs x(t) into a basal dendritic compartment b(t). Somatic activation s(t) is a combination of basal activity b(t) and recurrent apical feedback a(t). **e)** Schematic of the full model, including action neurons, which receive input from the representations in the CA1 network and dictate the agent’s decisions in its sensory environment, providing a closed loop between the environment and the network.

This is one example of a more general problem: real-world tasks often depend on complex combinations of sensory information, internal states, context, etc. which themselves are unknown prior to learning. Agents in these tasks must either pre-assign or learn state representations to create an actionable map of the environment. In animals, this latent structure is often referred to as a “cognitive map”, the idea of which has been very closely tied to the observed activity of hippocampus^6–8^. Cells in CA1 display fields specific to a variety of environment and task variables, including place, time, lap, evidence accumulation, and more^9–15^. These fields can emerge over the course of task-learning and can be induced artificially at the single cell level^16–18^. If activity in CA1 is indeed consistent with a cognitive map, how might biological agents perform representation learning in order to form these maps?

First, one could imagine a network of neurons which has pre-existing complex representations prior to the task. For example, one might consider using a temporal buffer in the network, or feeding activity into a liquid state machine, so that all possible sequences create a separate “latent” state^19^ (**Figure 1b**). However, reinforcement learning in these overcomplete state spaces is very slow and computationally expensive, scaling exponentially with the number of states (the so-called “curse of dimensionality”). A second option is to learn the structure of the environment through prediction and inference^19^ (**Figure 1c**). Some recent example models of the cognitive map include the Tolman-Eichenbaum Machine (TEM) and the Clone Structured Cognitive Graph (CSCG)^20,21^. The CSCG for example, presumes that each state initially has many copies, and then uses an expectation maximization (EM) algorithm to modify connections between these copies. However, the number of copies chosen for each state is still a parameter chosen a priori by the modeler. Further, plasticity rules involved in these inference models are generally non-local, rendering them difficult to map onto specific biological processes in hippocampus.

To theorize how hippocampus might form cognitive maps in a biologically plausible manner, we propose a model in which hippocampal CA1 uses local, single-cell plateau-based learning rules to develop population-level cognitive maps, allowing it to learn complex tasks. Notably, our model learns these state spaces and tasks simultaneously in an iterative manner. Cells encoding abstract state variables arise via learning the abstract logic of a given task, rather than existing a priori. Our model’s results are compared and validated against recent experimental results, in which the induction of splitter cells was only possible when a representational “split” was required to solve the task^18^. Finally, our model makes testable predictions about the potential codependence between latent hippocampal representations and behavior.

## Results

### Feedback in three-compartment CA1 neurons Enables Context-Sensitive Representations

To demonstrate how CA1 might learn task-dependent representations mechanistically, we create a three-stage closed-loop model. The first stage consists of the external environment, which is a Y-maze traversed by our agent in 2D Euclidean space. The second stage is a network of model CA1 neurons, which receive both external and recurrent inputs. The activity of the CA1 network is projected to a third stage, a set of “action” neurons which dictate the agent’s decisions within the external environment. As the agent learns a given task, the CA1 network develops appropriate latent representations upon which the agent can develop a successful policy.

Each neuron in our model CA1 network has three compartments – the soma, the apical dendrites, and the basal dendrites (**Figure 1d**). The basal dendrites receive input about external sensory information, while the apical dendrites contain feedback coming from a recurrent CA1 somatic - EC - CA1 apical dendritic feedback loop. For simplicity, we describe this feedback loop mathematically via a single recurrent weight matrix, W_ij_ (see Methods). The soma receives input from both dendritic compartments, such that the somatic activity s(t) outputs a combination of external information (basal activity, b(t)), modulated by the recurrent feedback which encodes the latent structure (apical activity, a(t)). The degree to which the somatic activity is dependent on recurrent, apical feedback is determined by a modulatory factor which depends on the sum of the incoming synaptic weights onto the apical dendrites (see Methods). In practice, this modulation means the soma essentially reflects external inputs in the absence of strong apical weights, and in the presence of strong apical weights, it reflects the co-tuning of external basal inputs and recurrent apical inputs. While we do not directly model a biophysical process for this compartment-specific modulation, the resulting increased co-tuning of the apical compartment with the soma can be compared to recent experimental findings of the same effect following manipulations of intracellular calcium release in vivo^22^.

Learning in the model consists of a plateau-dependent three-factor rule (see Methods), in agreement with experimental results which observe the formation of CA1 fields after so-called “plateau potentials”^23–25^. This rule depends on temporally filtered pre-synaptic activity, post-synaptic activity, and reward above expectation, and is triggered upon the occurrence of a plateau potential at time t_*plateau*_. These plateau events update the recurrent weights W_ij_, which determine the modulatory recurrent feedback housed in a(t). Crucially, this rule allows the network to reorganize sequentially activated pairs of states that lead to reward into a new “state”. In other words, this allows first-order states s_first_ = x_i_ (e.g., place only) to be combined into second order states s_second_ = x_i_x_j_ (e.g., place and cue context). Higher order states can then be learned as combinations of first and second order states, etc. Simpler learning rules which only consider one state and its connection to reward (i.e., TD) seem therefore incapable of learning these higher-order combinations, and more complicated rules which can learn higher order relations such as backpropagation within a recurrent neural network are typically non-local and therefore not typically biologically plausible^26^. Further, the inclusion of reinforcement above expectation (a “reward prediction error” of sorts) in our rule is in line with experimental results which show that dopamine receptors are necessary for CA1 to flexibly track changes to external environments and tasks^27–29^.

Attached to our hippocampal network is a network of two “action” neurons (representing the two possible turn directions, left or right) which dictate the agent’s turning decision in the environment. These action neurons v(t) are connected to somatic activity s(t) through weights Q, which can be interpreted as state-action values. The information in the model thereby follows a closed loop: The agent moves in the environment at a constant velocity ***dx***/***dt*** and observed external stimuli x(t) are fed into our representation layer as basal activity b(t). The basal activity b(t) combines with recurrent feedback a(t) to produce internal representational states s(t), which then activate action neurons v(t) through weight matrix Q. If the agent is at the choice point, these action neurons will dictate it to turn right or left, otherwise it will continue along its course before the process repeats (**Figure 1e**).

### Network learns task-dependent latent representations and does not integrate task-irrelevant representations

To test our model, we place an agent in a Y-maze environment to mimic that used in a recent experiment which concerned the emergence of splitters^18^. The agent is presented with one of two possible visual cues, A (red vertical bars) or B (black horizontal bars), before walking along a track (C). Upon reaching the end of the track, the agent can turn left or right (D or E) to potentially receive a reward (**Figure 2a**). We train the agent on one of two possible tasks. For the first task, reward was randomly given, such that D or E had a 50% probability of containing reward, regardless of the visual cue shown (random reward task, **Figure 2c**). In another task, the reward was contingent on the initial cue which was presented to the agent, such that if the agent saw cue A, the reward would be in port D with 100% certainty, and if the agent saw cue B, the reward would be in port E with 100% certainty (cue-dependent task, **Figure 2d**). In both tasks, we use the same induction protocol, wherein half of neurons receive location-specific plateaus after presentation of A, and the other half of neurons receive location-specific plateaus after presentation of B (**Figure 2b**).

**Figure 2.**
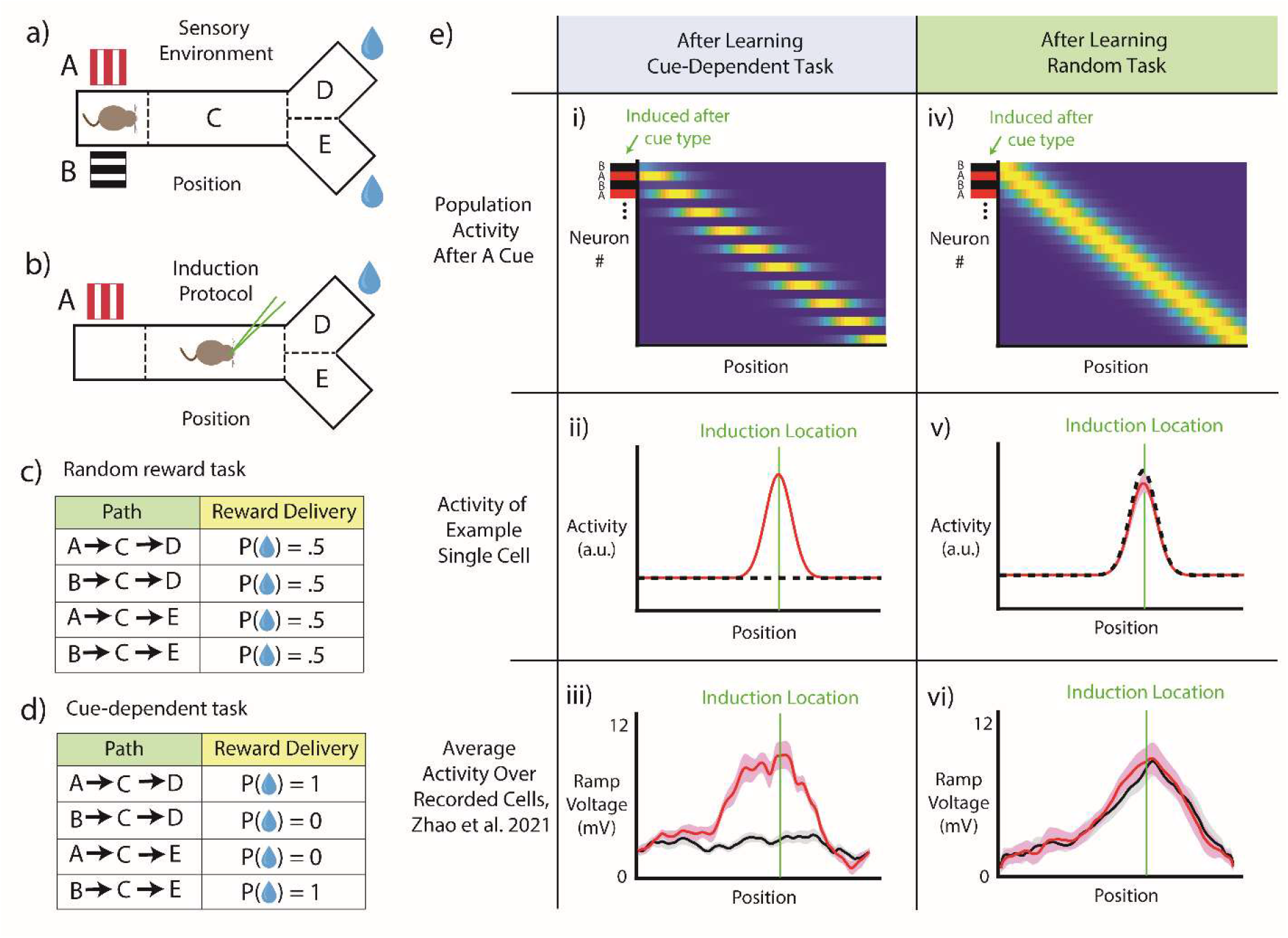
Splitters Emerge When Dictated by Task Structure. **a)** Task environment for simulation (to mimic the experimental design of Zhao et al.^18^). The agent is presented with one of two possible cues, A (red vertical bars) or B (black horizontal bars), before advancing through the track (C). It can then choose to go either D or E to potentially receive a reward. **b)** Fields are induced in hippocampal CA1 cells via the induction of a “plateau potential” at a certain location after a certain cue (in this case, A). This is designed to induce “splitters”, or place fields which only fire after the A cue. **c)** The random reward paradigm, where locations D and E each result in a 50% chance of reward delivery, regardless of the cue shown. **d)** The cue-dependent paradigm, where location D contains reward if preceded by cue A, and location E contains reward if preceded by cue B. **e)** Simulated population activity (i,iv), single cell activity (ii,v), and experimentally recorded activity (data adapted from Zhao et al.^18^) (ii,vi), for both the cue-dependent task (i-iii) and the random reward task (iv-vi). Notably, the same induction protocol generates splitters in the cue-dependent case, and non-specific place fields in the random reward case.

We observe that, in the cue-dependent task, neurons in our model CA1 develop “split” representations in agreement with the logic of the induction protocol. That is, cells which were artificially injected with current at a given cue-location combination during training develop fields that are selective to that cue-location combination after training (**Figure 2e,i**). An example cell (**Figure 2e,ii**) only fires at position C following presentation of the A but has no firing field at any location after the presentation of B. This result can be understood if we consider the dependence of our learning rule on reward, *dW* ∝ s_post_ ∗ s_pre_ ∗ (*r* − r_O_). State pairs which lead to increased reward in the task are potentiated within the recurrent weights, and thereby the activity of apical dendritic compartment a_post_(*t*) will depend on state s_pre_. Let’s examine the case where the reward location depends on the cue. The state pair s_D_ ∗ s_A_ will always lead to reward, and thereby the state s_D_ = *D* will become s_D,_ = *D* ∗ s_A_. Once our state representation at D becomes dependent on the cue, we can look one step backwards, as now the pair s_C_ ∗ s_D’_ = s_C_ ∗ *D* ∗ s_A_ will always lead to reward. Owing to this, *W*_CA_ will potentiate and s_C_ will also become the cue-dependent s_C,_, and the process repeats. The development of cue-dependent splitter cells we observe in the cue-dependent task is in agreement with recent experimental findings of task-dependent hippocampal representations^9,15,18,30–33^ (**Figure 2e,iii**). The obvious implication is that CA1 learns at least some of its representations along with the task, inconsistent with the assumption that all complex state representations exist a priori (**Figure 1b**).

However, if we train the agent on the random reward task, we observe that even when we induce plasticity only for a given cue-location pair, the resulting fields do not retain this information, instead becoming generic “place” fields (**Figure 2e,iv**). The same example cell which had in the cue-dependent task had developed a conjunctive field (location C if preceded by A), in this case encodes generally for location C, regardless of the preceding cue (**Figure 2e,v**). If we follow the same logic as before, our learning rule’s dependence on reward also predicts this result. The lack of splitting occurs because the reward is not exceeding expectation, since there are no state pairs in the random task which lead to reward above chance. As a result, the recurrent weights do not evolve, and the state representations do not pick up cue-dependence. Experimentally too, a cue-dependent field could not be induced in a single cell when the animal was trained on the random task^18^(**Figure 2e,vi**). In this scenario, an artificial agent would not require cue-dependent state spaces to form a good policy.

The more nuanced implication of these results is that a single postsynaptic cell may not have access to a complete pool of presynaptic inputs to form complex task representations in plateau-like plasticity, prior to task learning. This idea acts in contrast to reservoir type models, where an output has access to a pool of input which form a complete basis set, and few-shot learning can quickly “select” any desired representation of the inputs, without needing to train beforehand on any task. Taken together, our simulations suggest that both task and representation can be learned simultaneously, as opposed to complex task representations all existing, or being accessible, a priori. At the single cell level, this may suggest that complex post-synaptic representations depend on network-level rewiring of the *available* pre-synaptic connections during behavior, such that these connections are eligible for potentiation during events of plateau-driven plasticity (**Supplemental Figure 1**).

### Behavior co-evolves with internal representation of environment

From the simulations shown above, we can observe the state representations and behavior before and after learning (**Figure 3a**). However, since both internal state representations and behavior are plastic, we can also examine their dynamic evolution *during* learning. We quantify behavioral performance by measuring a running average of the fraction of correct turns the agent makes during training on the task. To quantify the state representation, we introduce a measure of “splitness” for neurons in our CA1 network (firing rate on A cue trials – firing rate on B cue trials). Since the task requires split representations to be solved, this can be understood as a sort of “fitness” of the state space to the task.

**Figure 3.**
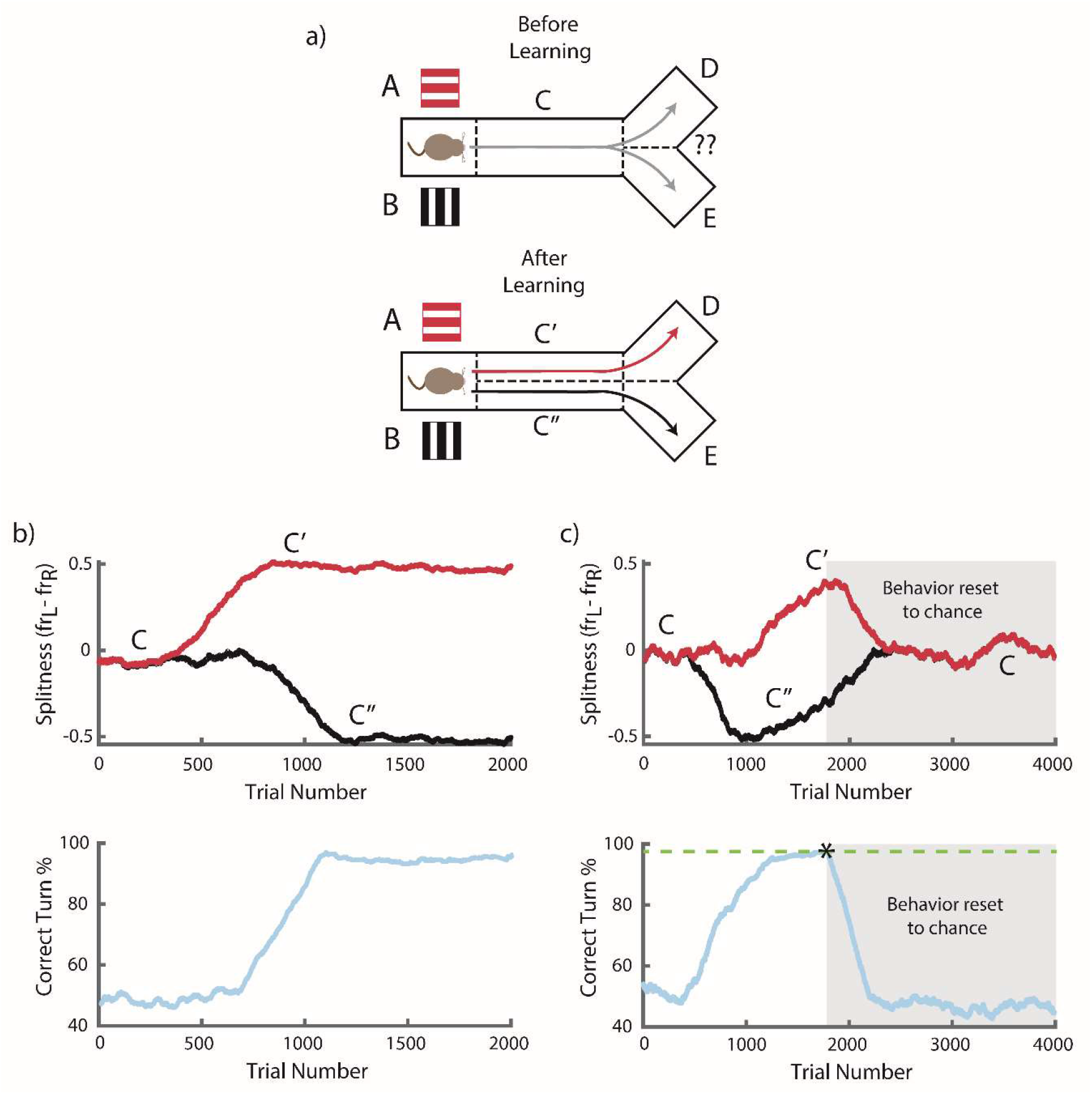
Behavior and Representation Iteratively Improve Each Other. **a)** Schematic of agent behavior before (top) and after (bottom) learning. Initially, the agent cannot implement the optimal policy at C since a single state cannot incorporate two separate state-action values. After training, the state has been split, and the policy has been learned. **b**,**c)** Top, difference in firing rates on A and B cue trials (“splitness”) of two example neurons over the course of learning. Bottom, behavioral performance, shown as the percentage of correct turns over the course of learning. **b)** Agent learns to maximize reward in the task over 2000 trials, with the standard plasticity protocol. Notably, for this simulation, the time course of the A and B representations is not aligned, with A splitters emerging before B splitters. Behavioral improvement begins concurrently with the existence of both A and B splitters, both of which are required to solve the task. **c)** Behavior is set back to chance once the agent reaches 95% task performance (green dotted line), chance behavior indicated by grey boxed area. Representation re-merges since neither C’ nor C’’ results in positive reward above expectation. In this simulation, B splitters emerge before A splitters.

We find that improvement in behavior of the agent in our model is directly tied to improvement in the “splitness” of representations (**Figure 3b**). In particular, the agent does not learn the optimal policy until both types of splitters, i.e. A-type and B-type, exist in the latent representation. For the particular simulation shown in (**Figure 3b**), A-type splitters form first, around trial 500, and B-type splitters follow around trial 750. Behavioral performance improves alongside the emergence of B-type splitters. From the perspective of reinforcement learning, learning of appropriate actions is contingent on an accurate state space, so one might expect the evolution of behavior to lag the evolution of state representations (τ_behavior_ >= τ_state_). However, here behavior and representation evolve *together*, (τ_behavior_ ∼= τ_state_) because 1) we cannot learn the split behavior without splitters, and 2) *we also cannot learn the splitters without split behavior*. Our model is able to break this loop owing to stochasticity in behavior, which acts as a symmetry-breaker to the underlying representation. Our learning rule reinforces breaks along dimensions useful to the task, while breaks along null dimensions will relax back to zero. Improper policy (i.e. going to the wrong reward port) will degrade the state representation, as unrewarded state pairs are depressed. Meanwhile, proper policy reinforces state pairs which led to reward. Therefore, in our model, proper state representations do not exist in the absence of behavioral performance, and vice-versa. These results are in agreement with experiments which observe task-relevant hippocampal representations arising on the same timescale as the relevant behavior^9,30,31,33^.

Experiments have also found that task-relevant representations are more stable than generic fields^30,31^, which may underlie their importance in supporting and maintaining task-performance. To test this codependency between state representation and task performance in our model, we performed a perturbation simulation where we reset behavior to chance once the agent has learned the task. In that case, the state representation in our model re-merges, since neither C’ nor C’’ results in positive reward above expectation (**Figure 3c**). This is in agreement with our earlier results in the random reward task (which is functionally equivalent in this case). We also perturb the network by selectively ablating splitters after training, which leads to a behavioral deficit and a degradation in the remaining latent representation, even in cells which were not removed (**Supplemental Figure 2**). These results could suggest that the disappearance, relabeling or remapping of fields could be functionally connected to a lack of relevance to behavior.

### Biases in representation lead to biases in behavior and vice versa

One consequence of the dynamic link between representation and behavior in our model is that biases in latent representation can lead to biases in behavior, and vice versa. To demonstrate this more concretely in our model, we first manually induce a bias in our network’s latent representation (**Figure 4a**). To do so, in half of our neurons we restrict our plasticity protocol to only induce fields after presentation of cue A. The remaining neurons are induced uniformly across cues. Measuring the splitness shows that this produces two functional populations: A-dependent splitters (C’) and generic place cells (C) (**Figure 4b**). We then examine how behavior co-evolves with this biased representation. We observe that on A trials, the agent always learns to always turn to D (correct policy), while on B trials, the agents on average behave at chance (**Figure 4c**). This occurs even though no restrictions are placed on behavioral learning. Instead, since the latent representation lacks B-specific splitters, the agent cannot assign unique state-action values in the B trials, and therefore fails to find the correct policy following the B cue.

**Figure 4.**
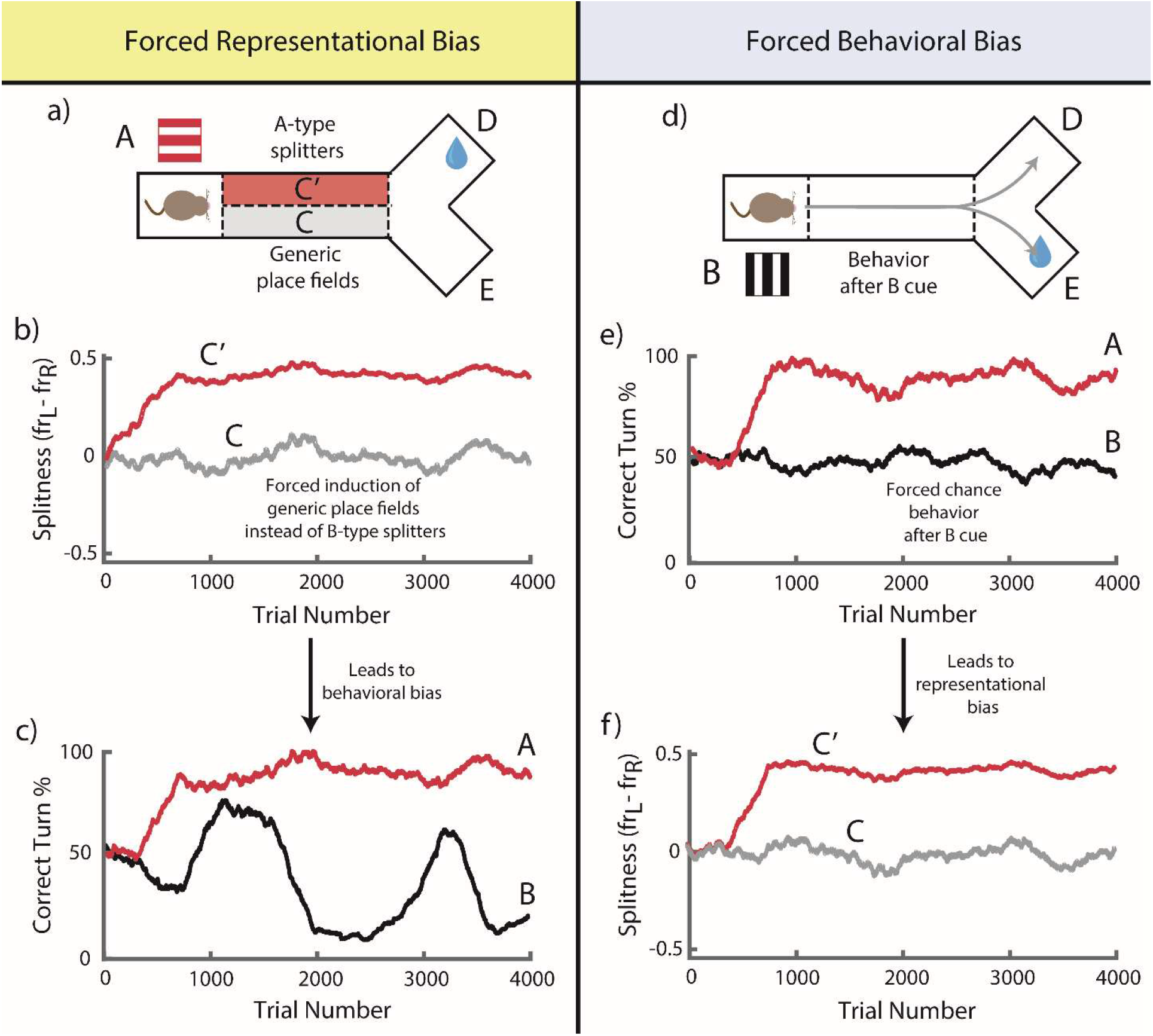
Biases in Representation Lead to Biases in Behavior and Vice Versa. **a)** Schematic of biased representations after presentation of A (top) and after presentation of B (bottom). Bias is forced upon the state space by manual induction general place fields (C) and A-dependent splitters (C’), but not B-dependent splitters. **b)** Difference in firing rates on A and B trials (“splitness”) of two example neurons over 4000 trials of learning. A non-specific place field (corresponding to C) is shown plotted in grey, and an A specific field (corresponding to C’) is plotted in red. **c)** Behavior, plotted as the fraction of correct turns on A trials (red) and B trials (black) over 4000 trials of learning. The agent cannot learn the optimal policy on B trials since the representation lacks B-dependent splitters. **d)** Schematic of forced behavioral bias after presentation of A (top) and after presentation of B (bottom). Behavior following presentation of A is allowed to learn as normal, but behavior following presentation of B is restricted to select one of D or E at chance. **e**,**f)** Same as in panels b,c), but for the case of forced behavioral bias (e) leading to representational bias (f). Notably, either a forced bias in induction protocol (panels a-c) or a forced bias in behavior (panels d-f) produce symmetric biases in both behavior and representation.

In a fixed-state RL model with an appropriately defined state space, suboptimal behavior generally arises from suboptimal value-learning. In contrast, what our model demonstrates here is *feature-specific* suboptimal behavior (i.e., only following the B cue), which is inconsistent with a general failure in value learning, and instead arises from suboptimal *representation learning*. In turn, our model predicts that feature-specific or abstraction-specific biases in behavior are the direct result of feature-specific or abstraction-specific biases in the underlying state representation in hippocampus. One way to test this experimentally would be via the selective ablation of feature-specific splitters, which in our model leads to a similar “biased” behavioral deficit as described above (**Supplemental Figure 3**).

To illustrate the inverse dependency, we allow our latent representation to learn as normal, but force behavior to operate at chance on B cue trials (**Figure 4d**). As expected, the agent successfully learns to choose the D port following the A cue, as we only restrict behavioral learning in the B cue trials (**Figure 4e**). Measuring the splitness of the resulting representation, we observe a striking symmetry to the case where we restricted our plasticity protocol directly, as the network again develops the same two functional populations: A-dependent splitters (C’) and generic place cells (C) (**Figure 4f**). This shows that not only can biases in representation lead to biases in behavior, but also that biases in the behavior of our agent can lead to biases in our model’s latent representation. A recent experimental result has shown that the quality of hippocampal place field maps is degraded when animals disengage from a virtual navigation task^34^, which supports the idea that hippocampal representations are, at least in part, shaped by behavior. Our model would predict that this result can be extended further, and that behavior and representations can be bidirectionally perturbed such that degradations in one will lead to degradations in the other, and improvements in one will lead to improvements in the other.

## Discussion

The hippocampus has long been thought to operate as a “cognitive map”, but the process by which it forms these maps is still unknown^6–8^. A principal difficulty in building cognitive maps is learning the appropriate state representation for tasks or environments which are described by complex, higher-order relations. Various models have been introduced to learn artificial state spaces (state discovery/representation learning)^35–37^, but our model attempts to directly link the emergence of population-wide cognitive maps to observed single-cell, plateau-based plasticity mechanisms in hippocampus. Further, our model replicates recent experimental results which show CA1 cells forming complex fields only when they are relevant to a behavioral task^18^. We observe in our model that behavior and representation iteratively improve each other, and that controlled modifications to either can lead to changes in the other.

Our model network learns to behave by learning the task structure directly, rather than building a general map of an environment. While it is true that hippocampal representations emerge in environments without an explicit task structure^12,13^, and common representations (such as physical space) are reasonable for an agent to generalize and learn in an unsupervised manner, unsupervised learning of higher-order state “environments” quickly becomes untenable, as the number of potential higher-order states grows exponentially with the size of the first-order state space (curse of dimensionality)^2,3^. As such, it is likely useful to learn only those higher order, complex representations which are necessary to maximize objectives related to explicit reinforcement or self-supervision. Evidence shows that complex representations in hippocampus are either a) more likely to, or b) exclusively emerge in tasks which require them^9,15,18,30,32^. Further, even representations of physical space have been shown to degrade with reduced task attention^34^, so perhaps even place fields can also be understood as a type of “task-related” representation, with the caveat that the task of locomotion nearly ubiquitous in any sort of natural behavior.

Our model considers fixed plasticity protocols, i.e., we as an outside observer choose to induce splitters in this task. While we do show that induction protocols unnecessary for the task are not integrated into the network, our network does not spontaneously generate plateaus. In natural formation of fields in CA1, of course, this loop would also be closed, and likely depends on other areas such as the entorhinal cortex^38^. One can imagine that through a noisy process, the network might sample potential latent states via one-shot learning to create quick combinations of external inputs/recurrent feedback. If these latent states are useful, perhaps they are reinforced and remain in the state space, remapping otherwise. It remains to be seen how well randomized or self-generated induction of plasticity could be implemented in our model.

It is worth noting that most high-order task logic will have many redundant solutions in lower-order spaces. For example, in the cue-dependent task we show here (**Figure 2d**), cue orientation and cue color are both predictive of reward location. So, either (or a combination of both) could equivalently influence the cognitive state. The resulting latent sequence would be identical, but the environmental stimuli which underlie it may differ, and perturbations to the environment (translocations, cue removal or replacement, etc.) may have confounding effects on the agent depending on which redundant solution to the task it has found, and how it maps onto sensory stimuli. Our model avoids this confound by predicting a direct equivalence between the structure of behavior and the structure of latent state representations. In this work, we show that the ad-hoc measure of representational “splitness” corresponds to behavioral performance (correct turn %) in our model, but more generally one might consider that any low-dimensional “task-relevant” measure of the representation may evolve in tandem with behavior (for example, increasing orthogonality of task-specific maps corresponding with improved performance in a discrimination task^30^).

Our model hippocampal network presented here demonstrates that single cell plateau-based learning can interact with behavioral learning to generate a population level cognitive map via cooperative improvement of behavior and representation. This method avoids problematic a priori assumptions of the structure of a given state space and presents a potential pathway for further research into the online, in vivo formation of task-relevant hippocampal cognitive maps.

## Methods

### External Input

The external sensory environment is modeled in the form of stereotypical 1-D positional tuning curves of the following form:

**Equation 1 – Positional Input**

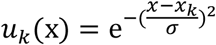

where k indexes over K total inputs, and *x*_k_ are the locations of the tuning curve centers, which have standard deviation σ. For simplicity, we model the animal as running at constant unit velocity through the track, such that *x* = *t*. CA1 neuron *i* receives input *M*_ik_*u*_k_(*t*). *M*_ik_ is equivalent to the identity matrix in this work, though in general, one might consider these weights to be plastic.

### CA1 Network

The rate activity of each CA1 neuron *i* is described by the following three-compartment model:

**Equation 2 – Somatic Activity**

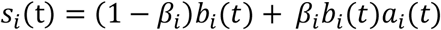

**Equation 3 – Contextual Factor**

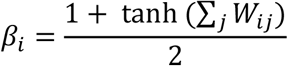

**Equation 4 – Basal Dendritic Activity**

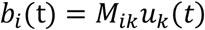

**Equation 5 – Apical Dendritic Activity**

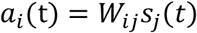

The neuron receives external input through basal dendritic activity *b*_i_(t) and internal recurrence through apical dendritic activity *a*_i_(t). The somatic activity is comprised of two components: basal activity *b*_i_(t) and basal-apical product *b*_i_(t)*a*_i_(t). The degree to which the somatic activity is influenced by each component is modulated via a non-linear sum of weights *β*_i_. In short, when the incoming weights to the apical compartments is large, the somatic activity is approximated by *b*_i_(t)*a*_i_(t), while when the incoming apical weights are small, the somatic activity is essentially *b*_i_(t). The soma’s dynamic sensitivity to apical activity is not based on one particular biophysical process, but might be similar to recent experimental findings of the increased apical co-tuning with the soma following manipulations of intracellular calcium release in vivo^22^. Recurrent weights W_ij_ are modeled here as a single matrix connecting CA1 somata to other CA1 apical dendrites. Owing to the biophysical wiring of hippocampus, this matrix is likely better described as two separate matrices, one which connects CA1 to EC (*R*_kj_), and another that projects from EC back to the apical dendrites of CA1 (*P*_ik_). In our linear model, these two explanations are equivalent, as we can rewrite the apical activity as the following:

**Equation 6 – Apical Activity Equivalence**

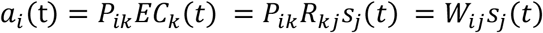

Where EC activity *EC*_k_(*t*) = *R*_kj_*s*_j_(*t*), and apical activity *a*_i_(t) = *P*_ik_*EC*_k_(*t*), leading to the relation describing in terms of the “biophysical” matrices *P*_ik_ & *R*_kj_:

**Equation 7 – Recurrent Weight Equivalence**

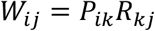

Somatic activity *s*_i_(t) produces eligibility traces *e*_i_(t):

**Equation 8 – Eligibility Trace of Somatic Activity**

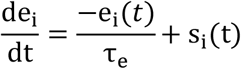

A history of activity for each neuron, e_i_ is calculated as an exponential filter of the spikes, with a time constant τ_e_.

**Equation 9 – Recurrent Three-factor Learning Rule**

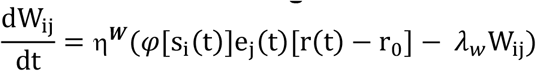

Recurrent weights W_ij_ are learned via a three-factor rule which depends on postsynaptic burst firing *φ*[s_i_(t)], presynaptic eligibility trace e_j_(*t*), and reward r(t) above expectation r_O_. Expectation r_O_ is defined as the average reward received by the agent, given a random policy. Burst firing occurs upon the induction of a plateau and acts as a pass-filter to the learning rule:

**Equation 10 – Plateau Firing**

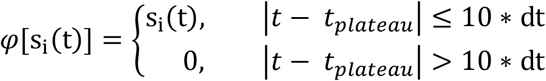

On unrewarded trials, weights of visited state pairs {i,j} will decay as *φ*[s_i_(t)]e_j_(t)r(t). However, for unvisited state pairs this term is zero, so weight decay − *λ*_w_W_ij_ is also included in the rule. Weights are updated in batch at the end of a given trial.

### Action Network

The main network is connected to an action network which determines the real space policy of the agent at the choice point. The activity v_l_ of units in the action network are described by the following:

**Equation 11 – Action Neuron Dynamics**

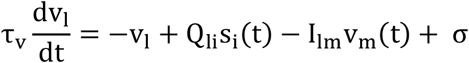

Where i indexes over the two possible choices for turn direction, left (L) and right (R). Q_li_ are the weights connecting the representation network to the action network, and I_lm_ provides mutual inhibition so that the action network has winner-take-all dynamics. Gaussian noise σ_N_ is introduced into the network, and the neurons operate with time constant τ_v_. The action network is queried at *t*_choic_ for the agent’s turn selection, which is decided by (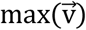), which results in the agent taking a turn either left or right into one of the reward ports.

**Equation 12 – Action Weight Learning**

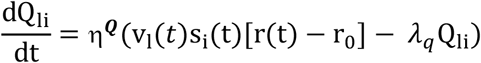

Action weights Q_li_ are learned via a three-factor rule, similar to that for Q_li_, except the action weights do not depend on presynaptic eligibility traces, so the three factors are presynaptic activity s_i_(t), postsynaptic activity v_l_(*t*), and reward r(t) above expectation r_O_.Though they do not explicitly match the definition of Q-values, they are effectively state-action values and thus can be interpreted similarly. As with the recurrent learning rule, weight decay − *λ*_q_Q_li_ is included so that unused state-action pairs depress over time.

### Network Architecture

The architecture of the network can be considered in three separate components: the external physical environment, a hippocampal latent representation network, and an action network. The external environment is a maze of length L, containing positions x, which are spanned by a population of N_inp_ generic tuning curves. Each tuning curve projects to the hippocampal network through the matrix *M*_ik_, which for this work is chosen to be identity matrix. As such, each neuron in the hippocampal network initially responds to a fixed location in real space, acting as a generic place field before training on a task. The hippocampal CA1 network is comprised of N neurons, each of which consists of three compartments: the soma, the basal dendrites, and the apical dendrites. The soma of presynaptic neuron j connects to the apical dendrites of postsynaptic neuron i via weight matrix *W*_ij_. The external environment feeds into the basal dendrites via matrix *M*_ik_. Somatic activity is determined by a combination of basal and apical activity (see **Equation 2**). The action network consists of two neurons representing the potential turn directions (R,L), and receives input from CA1 somata via the weight matrix Q_li_.The action network also contains recurrent inhibitory weights I_lm_. The agent’s decision in the environment to turn right or left at the choice point is determined by the action neuron with the highest activity at the time of the choice point.

## Model Parameters

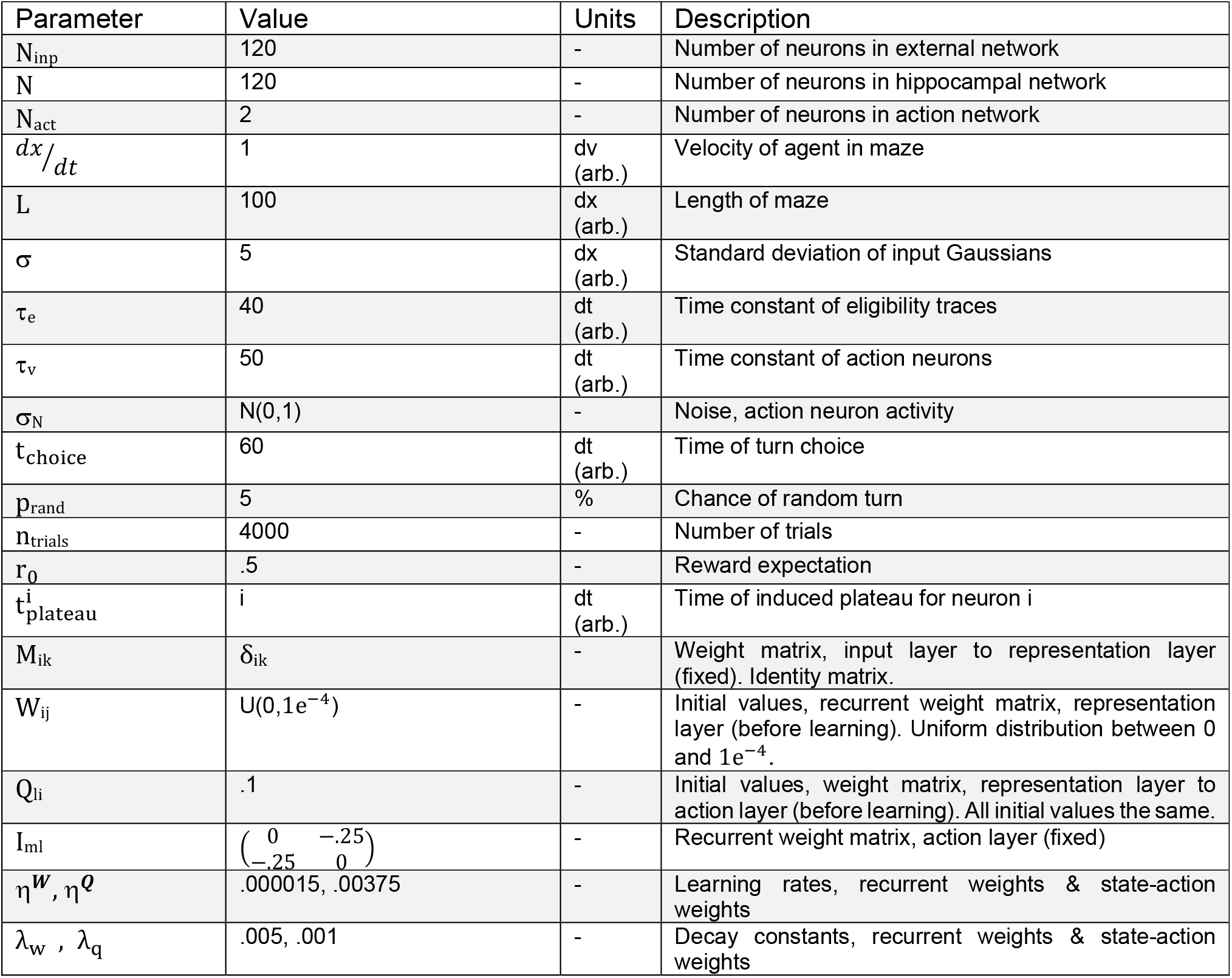

| Parameter | Value | Units | Description |
| --- | --- | --- | --- |
| $N_{\text{inp}}$ | 120 | - | Number of neurons in external network |
| $N$ | 120 | - | Number of neurons in hippocampal network |
| $N_{\text{act}}$ | 2 | - | Number of neurons in action network |
| $dx/dt$ | 1 | dv (arb.) | Velocity of agent in maze |
| $L$ | 100 | dx (arb.) | Length of maze |
| $\sigma$ | 5 | dx (arb.) | Standard deviation of input Gaussians |
| $\tau_e$ | 40 | dt (arb.) | Time constant of eligibility traces |
| $\tau_v$ | 50 | dt (arb.) | Time constant of action neurons |
| $\sigma_N$ | $N(0,1)$ | - | Noise, action neuron activity |
| $t_{\text{choice}}$ | 60 | dt (arb.) | Time of turn choice |
| $p_{\text{rand}}$ | 5 | % | Chance of random turn |
| $n_{\text{trials}}$ | 4000 | - | Number of trials |
| $r_0$ | .5 | - | Reward expectation |
| $t_{\text{plateau}}^i$ | i | dt (arb.) | Time of induced plateau for neuron i |
| $M_{ik}$ | $\delta_{ik}$ | - | Weight matrix, input layer to representation layer (fixed). Identity matrix. |
| $W_{ij}$ | $U(0, 1e^{-4})$ | - | Initial values, recurrent weight matrix, representation layer (before learning). Uniform distribution between 0 and $1e^{-4}$. |
| $Q_{li}$ | .1 | - | Initial values, weight matrix, representation layer to action layer (before learning). All initial values the same. |
| $I_{ml}$ | $\begin{pmatrix} 0 & -.25 \\ -.25 & 0 \end{pmatrix}$ | - | Recurrent weight matrix, action layer (fixed) |
| $\eta^w, \eta^q$ | .000015, .00375 | - | Learning rates, recurrent weights & state-action weights |
| $\lambda_w, \lambda_q$ | .005, .001 | - | Decay constants, recurrent weights & state-action weights |

## Acknowledgements

This work was supported by BBSRC (BB/N013956/1), Wellcome Trust (200790/Z/16/Z), the Simons Foundation (564408) and EPSRC(EP/R035806/1).

## Supplemental Figures

**Supplemental Figure 1.**
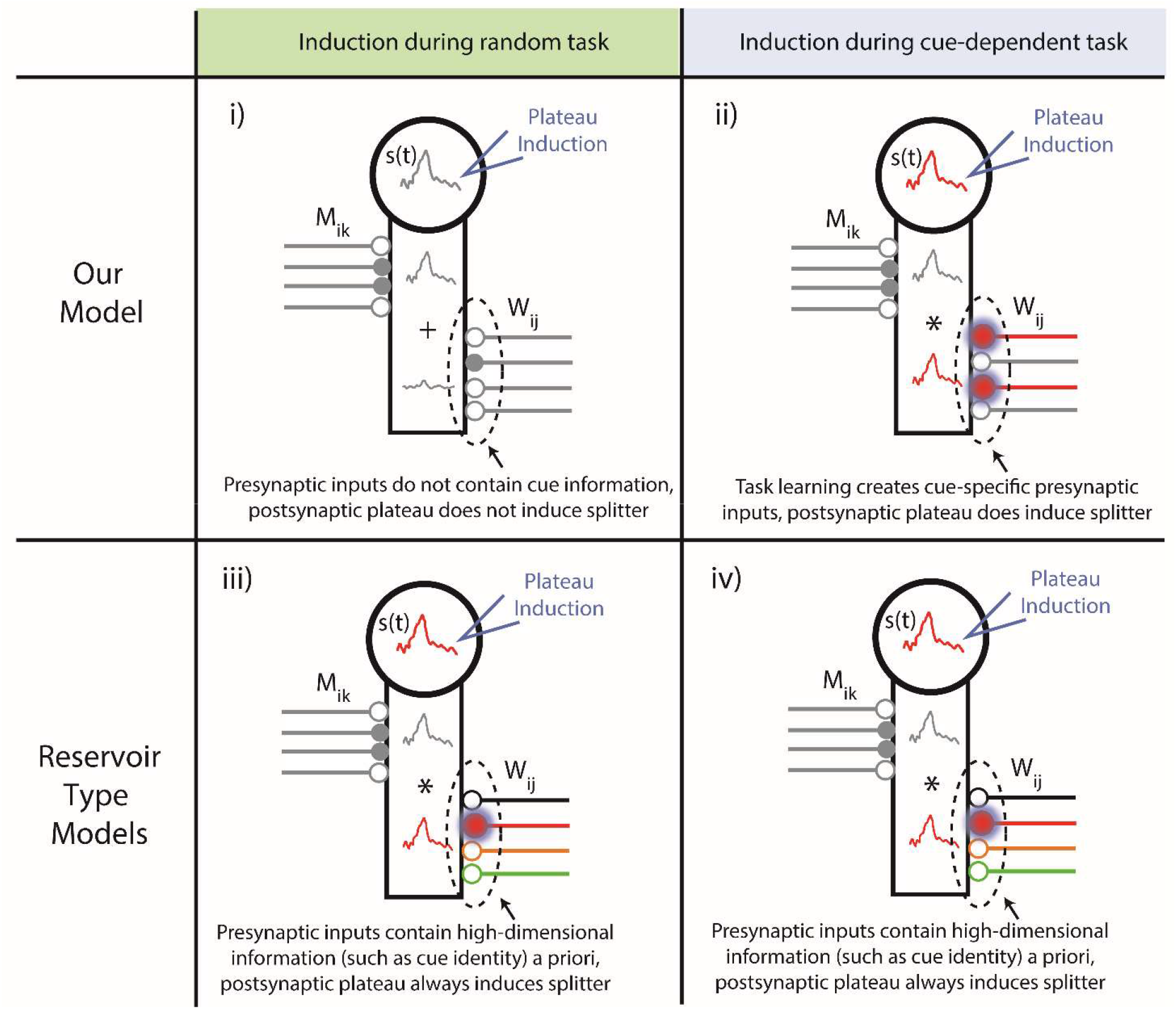
Single neurons require task-learning for few-shot induction of context-specific fields. Illustration of plateau-based induction at a single post-synaptic neuron for different model assumptions. Somatic activity s(t) is shown along with basal and apical dendritic activity. Grey inputs are non-specific, and red and black inputs correspond to the A and B cue contexts. Panels i & ii, our model, panels iii & iv, model with reservoir assumption. **i)** In our model in the random task, plateau induction is insufficient to induce splitters, since presynaptic inputs are initially non-specific and low-order (i.e. do not contain specific cue information). **ii)** Over the course of training on the cue-dependent task, the available pre-synaptic inputs reorganize to deliver higher-order (i.e., cue and place) information relevant to solving the task. Subsequent plateau induction can then trigger one- or few-shot induction of complex fields which are a conjunction of position and contextual information (in this case, the identity of the preceding cue). **iii, iv)** In models which assume the post-synaptic neuron has pre-existing access to many high-dimensional, abstract inputs (e.g., if we assume the postsynaptic neuron is the output of a large reservoir network), single shot plateau-based learning can immediately access the context which is selected for in the induction protocol, regardless of training on the task. Therefore, splitters arise in the reservoir model regardless of the task constraints, instead relying only on the logic of the induction protocol.

**Supplemental Figure 2.**
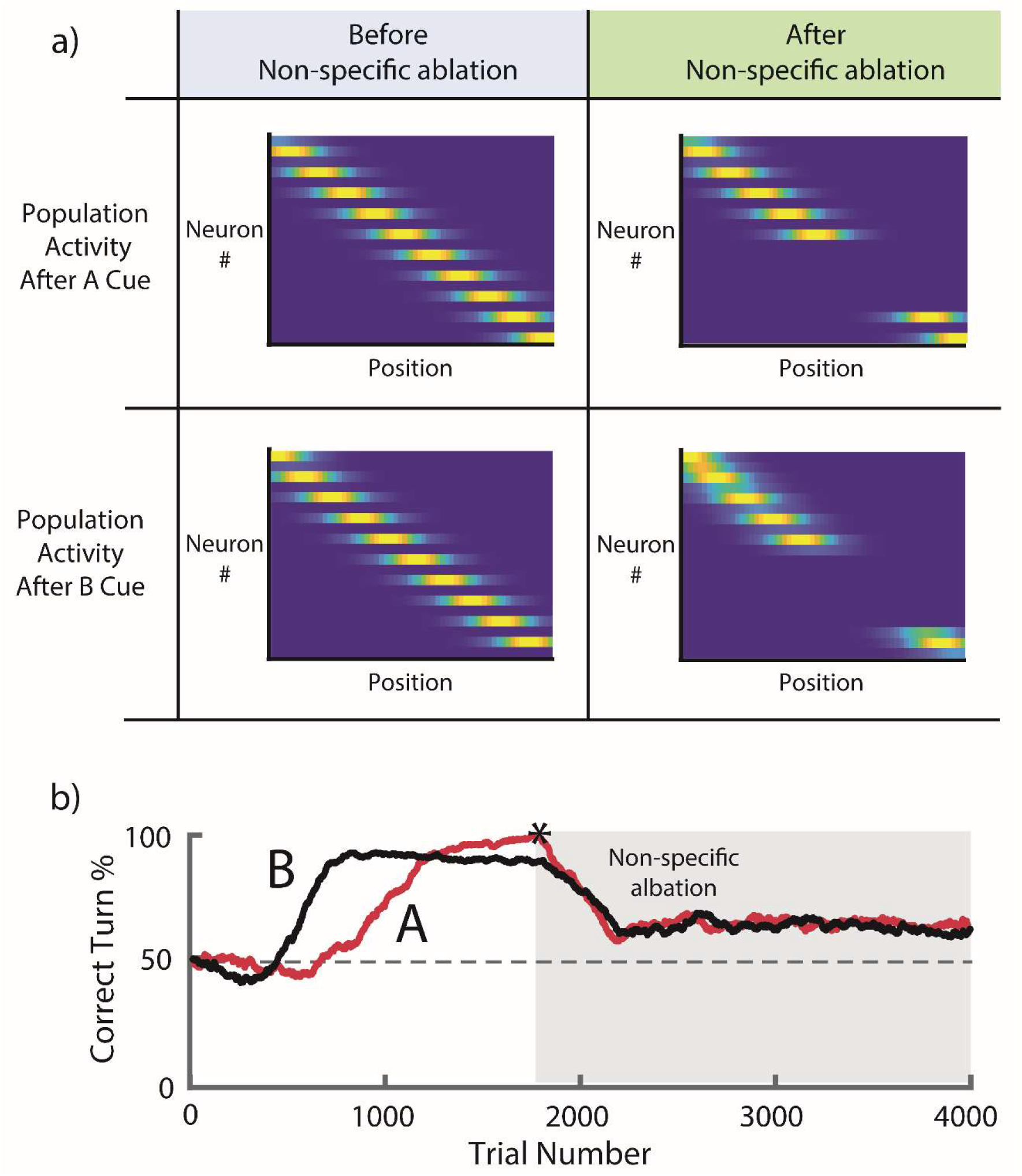
Non-specific ablation of splitter neurons negatively impacts general behavioral performance. **a)** A selection of both A-type and B-type splitters are ablated after behavior reaches criterion, leaving a gap in the representation. **b)** Behavior on A (red) and B (black) trials, plotted as the fraction of correct turns. The agent learns to maximize reward in the task until non-specific ablation (star), after which (grey box) performance degrades generally in both the A and B trials but remains above chance.

**Supplemental Figure 3.**
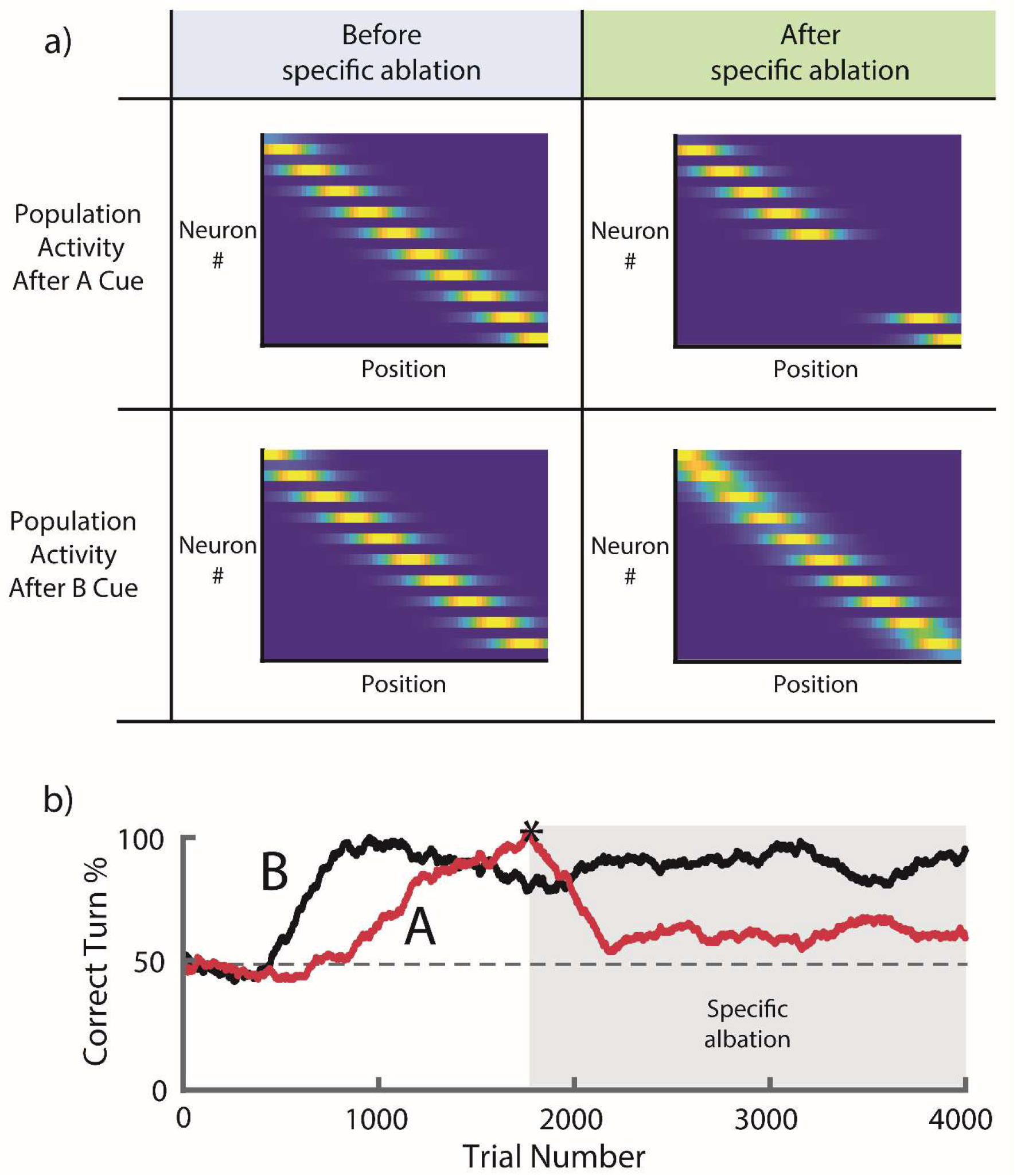
Feature-specific ablation of splitter neurons produces feature-specific behavioral deficits. **a)** A selection of A-type splitters is ablated after behavior reaches criterion, leaving a cue-specific gap in the representation. **b)** Behavior on A (red) and B (black) trials, plotted as the fraction of correct turns. The agent learns to maximize reward in the task until specific ablation (star), after which (grey box) performance degrades specifically in A trials, leading to a feature-specific behavioral deficit.

